# Global distribution of honeybee gut microbiome and pesticide-driven adaptations in opportunistic microbial species

**DOI:** 10.1101/2025.02.25.640077

**Authors:** Nelli Vardazaryan, Lusine Adunts, Inga Bazukyan, Magdalina Zakharyan, Honglian Liu, Chrats Melkonian, Lilit Nersisyan

## Abstract

Honeybees (*Apis mellifera*) rely on a specialized gut microbiome shaped by climate, flora, agrochemicals, and dietary supplements. Yet, how these factors alter microbiome composition and function remains unclear.

We integrated 16S rRNA and shotgun metagenomics data from eight published studies across six regions, alongside newly generated 16S data from Armenia, to assess how environmental and agrochemical factors influence the honeybee gut microbiome. We also introduced a novel co-abundance and functional analysis pipeline to identify treatment-affected bacterial networks and associated pathways.

We observed a stable core of six to twelve phylotypes in amplicon and metagenomic datasets respectively. However, we note significant geographic variation in relative abundances, likely reflecting differences in diet, climate, and local flora. Armenian data revealed distinct seasonal shifts, particularly elevated *Commensalibacter* in autumn and minor urban-rural differences. Pesticide treatments elicited varying responses: oxalic acid drove pronounced beta-diversity shifts; neonicotinoids had subtler effects, both primarily impacting opportunistic pathogens; and glyphosate disrupted core taxa with stronger effects in newly emerged bees under prolonged exposure. Co-abundance network analysis highlighted that the pesticide-associated community was enriched in adaptive pathways, including potential glyphosate degradation by *Pseudomonas*, biofilm formation, and aromatic amino acid synthesis.

These findings reaffirm the stability of the honeybee core microbiome yet underscore that environmental and anthropogenic stressors induce distinct compositional and functional shifts. We emphasize the need for longitudinal metagenomic approaches that enable high-resolution functional profiling and co-abundance network analysis to clarify how these microbiome shifts impact bee health and colony sustainability.

## Introduction

The Western honey bee (*Apis mellifera*) is a key pollinator, renowned for its broad geographic distribution and effective foraging [1,2]. In recent years, honeybee populations have declined globally, with annual losses reaching 30-45% [3–6]. Several factors have been linked to this decline, including climate change [6], heavy use of pesticides [7], use of antibiotics [8], and pathogen infection [9], but no single factor has yet been implicated as the cause.

The gut microbiome plays a vital role in bee health by regulating metabolism, pollen digestion, immunity, development, and behavior [10]. The core microbiome of adult workers comprises highly specialized bacteria. Five consistently reported phylotypes include Proteobacteria - *Gilliamella* (*G. apicola*) and *Snodgrassella* (*S. alvi*), which mostly reside in the ileum and form a biofilm; and Bacillota - *Bombilactobacillus* (*a.k.a Lactobacillus Firm-4*), *Lactobacillus* (*a.k.a. Lactobacillus Firm-5),* and *Bifidobacterium*, predominantly found in the rectum [11–14]. Other genera, such as *Frischella, Bartonella, Commensalibacter,* and *Bombella*, are also common [11,15,16], while *Apilactobacillus kunkeei, Apibacter, Serratia marcescens,* and some Enterobacteriaceae, are sometimes reported at low abundance [17].

Environmental factors, health status, and diet strongly affect the honeybee gut microbiome, often leading to increased mortality [4,18]. In particular, agricultural pesticides can disrupt the microbiota, altering behavior and immune function [19–21]. Neonicotinoid insecticides, such as thiacloprid, clothianidin, and imidacloprid, not only directly impair the nervous system affecting the memory and locomotion of honeybees, but may also disrupt the gut microbiome [22–25]. Oxalic acid, an acaricide insecticide commonly used against Varroa parasites and considered relatively safe for bees [26], can nonetheless lower the gut pH and alter the microbiome composition at high doses [19]. Glyphosate, the most widely used herbicide globally, negatively affects honeybee behavior and physiology [27]. In plants, it targets the EPSPS enzyme in the shikimate pathway, crucial for the synthesis of essential aromatic amino acids [14]. Since many bacteria possess glyphosate-sensitive class I EPSPS enzymes, concerns present about its adverse impact on the bee gut microbiome [14,28–30]. Among the glyphosate-affected species, *S. alvi* is most consistently reported. However, the overall impact of glyphosate on the gut microbiome remains inconclusive [4,29,30].

Although the honeybee gut microbiome is widely recognized as essential for bee health, conflicting evidence remains regarding how pesticides affect its taxonomic composition and functional potential. For example, studies on neonicotinoids yield contradictory results, some report negligible impacts under short-term or laboratory conditions, while others document significant shifts after long-term field exposure [25,31]. Similarly, while in vitro experiments show that oxalic acid increases the abundance of *Gilliamella, Bifidobacteria, Lactobacillus*, and *Snodgrassella*, these effects are not consistently observed in vivo [19]. The impact of glyphosate also varies considerably with dose, exposure duration, and the state of gut colonization [29,32,33]. Importantly, these studies do not fully explain the functional consequences or reveal the adaptation mechanisms underlying these taxonomic shifts.

To address these gaps, we performed a meta-analysis by aggregating honeybee gut microbiome datasets from eight published studies across six geographic regions and incorporated newly generated 16S data from Armenia, an understudied region in apicultural research. We employed a unified bioinformatics workflow to reprocess the data, thereby reducing technical variability and enabling robust comparisons of community structure and predicted functional capacity across diverse environments and pesticide treatments. Furthermore, we introduced a novel co-abundance network analysis to identify treatment-affected bacterial communities and elucidate their putative functional roles.

## Materials and methods

### Dataset search and inclusion criteria

We conducted a systematic search for datasets in the NCBI SRA database using the terms: *(microbiome AND “gut” AND (“Apis mellifera” OR “honeybee”))*. We included both 16S rRNA amplicon and whole-genome sequencing (WGS) datasets. Relevant datasets were additionally gathered from recent literature.

The inclusion criteria for studies were as follows: (i) paired-end sequencing; (ii) study subjects being worker bees; (iii) inclusion of the entire gut (including crop, midgut and hindgut); (iv) availability of raw data and corresponding metadata; (v) for 16s amplicon sequencing, libraries targeting the V4 region (e.g., V4 only, as well as V3-V4) to minimize batch effects related to variable amplicon region choice and enable cross-study comparison [34,35].

The data was downloaded from SRA using the *fastq-dump* tool from the NCBI SRA Toolkit, or downloaded directly through personal communication with authors. Detailed information about the datasets used is available in Table S1.

### DNA extraction and sequencing of the Armenian samples

Honeybees from different colonies were collected from the Gegharkunik, Kotayk, Syunik, Vayots Dzor, Lori, Ararat, Artashat, Jermuk, Hanqavan, and Aghavnadzor provinces of Armenia. Samples were retrieved from a single colony in each province to limit the effect of genetic variation. Guts were dissected using flame-sterilized forceps under aseptic conditions. The head was separated from the thorax to detach the esophagus, and the entire intestinal tract was carefully extracted through the anterior end by grasping the last abdominal segment with forceps and gently pulling. Additional segments of the abdominal exoskeleton were carefully detached to ensure the rectum was removed intact, as it was notably enlarged in most bees.

DNA was extracted using the cetyltrimethylammonium bromide (CTAB) bead-beating method, followed by phenol-chloroform purification and ethanol precipitation, as described in [36]. The extracted DNA was sent to Macrogen Inc. (Seoul, Korea) for paired-end 16S sequencing on the Illumina MiSeq platform. The V3-V4 hypervariable regions of the 16S rRNA gene were amplified using the primers Bakt_341F (5′-CCTACGGGNGGCWGCAG-3′) and Bakt_805R (5′-GACTACHVGGGTATCTAATCC-3′).

### 16S sequencing data processing

Raw paired-end reads were filtered out if they contained any of the following: (i) ambiguous bases, (ii) homopolymers longer than eight bp (using *fastq_qual_trimmer* v1.0), or (iii) quality scores less than 28 (using *fastq-mcf* v.1.0).

The trimmed sequence reads were processed using the *dada2* package for R (v.1.22.0) to infer exact amplicon sequence variants (ASVs) [37]. Briefly, reads were truncated with the *filterAndTrim* function, specifying the *truncLen* parameter to trim bases with a quality score below 30; reads containing more than two errors (forward) and three errors (reverse) were removed with the *maxEE* parameter; sequence reads were dereplicated, denoised, and merged using *dada2* default parameters, with pooled sample inference applied for each dataset.

The datasets were merged into a single ASV table using the *mergeSequenceTables* function of the *phyloseq* package (v.1.48) [38]. To reduce batch effects, ASVs were truncated to the common V4 region of the 16S rRNA gene through the following steps: (i) sequences were aligned using the *msa* R package v.1.30.1 [39]; (ii) aligned ASVs were truncated from the 3′ end, based on the C3 conserved sequence region (GTGCCAGCAGCCGCGGTAAA), allowing up to five possible mismatches; (iii) reads were further truncated at the 5′ end, to the most abundant length of 60 bp; (iv) alignment gaps were removed.

Taxonomy was assigned to the ASVs using the SILVA database [40] via the *assignTaxonomy* function in *dada2*. The taxonomy-assigned data was converted into a *phyloseq* object. Chimeric ASVs and those observed in only one sample were filtered out. ASVs unclassified at the genus level, assigned to mitochondria, chloroplasts, or eukaryotes, or not assigned to bacterial or viral lineages were excluded. To improve taxonomic accuracy, we retained only taxa meeting the following criteria: (i) at least 10 ASVs in one or more samples, and (ii) present in more than 70% of samples in each dataset or treatment group within a study.

Rarefaction curves were constructed using the *rarefy* function from the *vegan* package (v.2.6.8) to assess the number of observed ASVs relative to library size. Each sample was rarified to 12,000 reads using the *rarefy_even_depth* function in *phyloseq* (<u>Fig. S1</u>).

### WGS data processing

Taxonomic classification of the WSG reads was performed with Kraken 2 (v.2.1.2) [41] with default settings and the NCBI Standard database (https://benlangmead.github.io/aws-indexes/k2, accessed on 17 May 2022), without removal of host sequences. 1-9% of reads were assigned to prokaryotes.

Species-level counts were re-estimated using Bracken (v.2.9). Downstream analysis of Bracken results was conducted using the *phyloseq* package [42]. The species abundance table was rarified to 800 000 reads (Fig. S1).

### Diversity analysis and statistics

Shannon alpha diversity indices were calculated using phyloseq. For community composition (multivariate) analyses, pairwise Bray-Curtis dissimilarity between samples was estimated, and principal coordinates were computed using the *dist_calc* and *ord_calc* functions in the microViz package v.0.12.4 [43]. PERMANOVA analysis was performed using the *adonis2* function in the *vegan* package v.2.6.8. The *envfit* function in *vegan* was used to fit environmental variables onto the ordination axes using multiple regression. The statistical significance of these relationships was determined using a permutation test. The fitted environmental variables were visualized as vectors, projected onto the ordination diagram using the *geom_segment* function in *ggplot2* (v.2 3.5.1).

### Differential abundance analysis

To identify species or genera associated with treatment, we performed differential abundance analysis with the ANCOM-BC2 v.2.6.0 [44]. Unlike other methods, ANCOM-BC2 assumes that, under the null hypothesis, the proportions of taxa in a sample remain constant across conditions, making it particularly effective in addressing the biases and covariances common in microbiome datasets. It also adjusts for biases derived from varying sequencing and extraction methods. While running ANCOM-BC2, we encountered an error related to object length mismatches that we corrected in our pipeline using the steps suggested by the community (https://github.com/FrederickHuangLin/ANCOM-BC/issues/231).

### Co-abundance network analysis

We developed a new methodology and an R package, *micronet*, for the construction and functional analysis of microbial taxon-taxon co-abundance networks. We first selected taxa with at least 0.05% and a prevalence of 20% or more in at least one treatment group (e.g. control, treatment, and, if applicable, experiment group). If some taxa within a genus were identified at the species level, the rest of the unidentified ASVs were aggregated and marked with the genus name and “other” prefix. For genera with no ASV identified at the species level, we aggregated those and marked them with the genus name (Table S10). We calculated Spearman correlation coefficients between the relative abundances of taxa within each treatment group in each study. To ensure robustness, we performed bootstrapping using the *co_occurrence* function of the phylosmith R package: in 200 iterations per treatment group, a random subset of samples was selected, and only taxa pairs with an absolute correlation coefficient >= 0.6 and FDR-adjusted p-value < 0.1 were retained. We kept taxon pairs that appeared in at least 80% of the iterations with a consistent correlation sign. These consistently correlated pairs were then merged across groups into a single co-abundance network. Nodes represented species or genera, and edges represented significant co-abundances, with edge weights defined as the median correlation coefficient across the groups where the correlation was observed. This pipeline is implemented in the *generate_network* function of the *micronet* package.

Next, we aimed to detect treatment-affected subnetworks. We computed log-fold changes (logFC) of taxon relative abundances between treatment and control groups with *ANCOM-BC2*. The absolute logFC values were discretized into node weights as follows: |logFC| > 1.5: weight 3; |logFC| between 1 and 1.5: weight 2; |logFC| between 0.5 and 1: weight 1; |logFC| < 0.5: weight -2; missing values: weight 0. We then extracted treatment-associated subnetworks using the maximum weight connected subgraph (MWCS) algorithm implemented in the *mwcsr* R package (v.0.1.9). MWCS identifies a connected group of nodes that maximizes the sum of their weights, representing the most relevant set of connected taxa. We have implemented this pipeline into the *subgraph_analysis* function of *micronet*.

Finally, the networks were visualized in Cytoscape v.3.9.0 using an edge-weighted spring-embedded layout, in which nodes attract or repel each other based on their correlation coefficients [45].

### Genome assembly and taxonomic characterization

Metagenome-assembled genomes (MAGs) were reconstructed from WGS datasets. The resulting MAGs and functional annotations were later used as proxies to enhance functional calls in 16S datasets (see below).

Raw library sizes ranged from 2.1×10^6^ to 1.0×10^7^ reads. Reads were trimmed and quality-controlled with *fastp* (v.0.23.2); those <100 bp, with >10% low-quality bases (phred quality <20), >5 ambiguous bases, or with low complexity (with repetitive or homogeneous patterns) were discarded, and Poly-G/X tails were trimmed. Host DNA was eliminated by aligning reads to the *Apis mellifera* genome (GCF_003254395.2), using Bowtie2 (v.1.3.1). Post-processing yielded 7.8×10^4^ to 2.2×10^6^ paired-end reads per sample.

Both WGS datasets were co-assembled with MEGAHIT (v.1.2.9), and contigs were binned using MetaBAT, CONCOCT, and MaxBin. The three bin sets were merged via MetaWRAP’s *bin_refinement* module (v.1.3 [46]). Bin quality was assessed with CheckM (v.1.0.18); only bins with > 80% completeness and <5% contamination were retained, yielding 15 bins. Finally, MAGs were taxonomically annotated using GTDB-Tk (v.2.4.0) with default parameters [47] (Table S2).

### Gene function annotation

We predicted microbial gene functions using PICRUSt2 (v.2.4.1) [48,49]. ASVs with species/genus level taxonomic assignments were used to annotate genes with KEGG Orthology (KO). We refined KO predictions using functional profiles from corresponding WGS-derived MAGs to overcome the limitations of high-level ASV taxonomic annotations. MAGs were annotated with eggNOG-mapper (v.2.1.12) [49]. For species with both MAG and ASV annotations, only KOs from MAGs were retained (Table S2).

### Pathway enrichment analysis

We analyzed the refined KO table to identify enriched KOs in treatment-affected microbial communities. For each subnetwork, we separated nodes into up- or down-regulated groups based on positive or negative logFC values. KO-related gene counts in these groups were compared against the entire network using a hypergeometric test in R’s *stats* package (v.4.0.2). We computed the total probability of observing the given count (with *dhyper*) or any higher count (with *phyper*, lower.tail = FALSE). KOs with p-values < 0.1 were deemed significantly enriched.

Next, we performed KEGG pathway overrepresentation analysis with *enrichKEGG* in the *ClusterProfiler* package (v.4.12.6) [50], using the full network’s KO distribution as a reference. Pathways with FDR-adjusted p-values ≤ 0.2 were considered enriched. Only pathways enriched among up-regulated nodes and not enriched in down-regulated nodes were retained.

## Results

### Building a cross-study dataset with batch effect correction

We collected, reprocessed, and reanalyzed raw data from eight published studies. We systematically searched the NCBI SRA database for 16S and WGS datasets from adult workers. These datasets comprise samples collected from hives or sterile lab conditions, with or without pesticide or sugar supplement treatments, and span diverse geographic regions. Additionally, we generated an in-house 16S rRNA dataset from the entire guts of worker honeybees collected across Armenia (Fig. 1).

**Figure 1.**
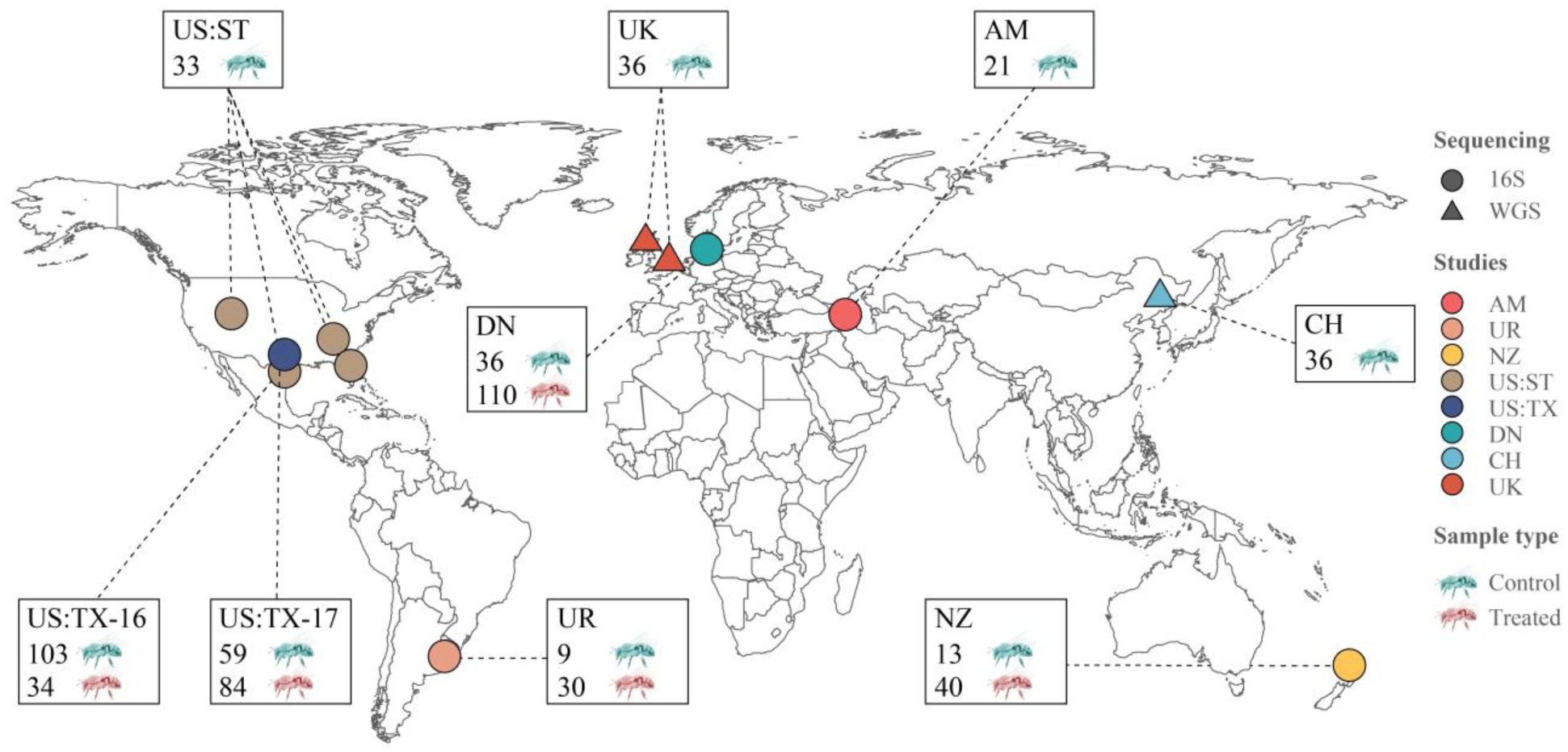
Geographic distribution and sample composition of studies included in the meta-analysis. The map shows study locations and the number of samples per treatment (red) and control (green) group. Colors indicate the sampling location: (AM: Armenia, DN: Denmark, NZ: New Zealand, US:ST: Southern US, CH: China, UK: United Kingdom). Shapes indicate whether the samples are from 16S rRNA or WGS sequencing.

In total, we analyzed 55 WGS and 461 16S samples. All WGS samples were untreated, whereas the 16S dataset comprised 190 untreated controls and 242 treated samples (Table 1). Treatments included herbicides (glyphosate), insecticides (oxalic acid and neonicotinoids), and sugar supplements. Mean library sizes varied widely, from 10K to 160K reads per library for 16S and WGS datasets, respectively. 57.2% of 16S samples targeted the V4 and 42.8% - the V3–V4 regions (Table 1, Fig. S1,and Table S1).

**Table 1.** Summary of dataset features included in this study.

| Dataset ID | Data description | Sample type<sup>A</sup> | Treatment | Bee source | Treatment/condition | # Samples |
| --- | --- | --- | --- | --- | --- | --- |
| AM | 16S, V3-V4 | Entire gut | Untreated | Apiary | Untreated | 21 |
| UR [4] | 16S, V3-V4 | Entire gut | Herbicide | Lab | Glyphosate<sup>B</sup> | 19 (10) |
| DM [19] | 16S, V4 | Entire gut | Pesticide | Apiary | Acetamiprid/Thiacloprid/Oxalic acid<sup>C</sup> | 182 (39/35/36) |
| US:TX-16 [25] | 16S, V4 | Crop-Rectum | Pesticide | Apiary | Imidacloprid<sup>B</sup> | 69 (34) |
| US:TX-17 [29] | 16S, V4 | Entire gut | Herbicide | Apiary/Lab | Glyphosate<sup>B</sup> | 84 (59) |
| US:ST [87] | 16S, V4 | Entire gut | Untreated | Apiary | Untreated | 44 |
| NZ [18] | 16S, V3-V4 | Crop-Rectum | Supplement | Apiary | Sugars (Dihydroxyacetone/Manuka honey/Invert sugar) or antibacterial (Methylglyoxal)<sup>C</sup> | 53 (8/16/9/8) |
| CH [88] | WGS | Entire gut | Untreated | Apiary | Untreated | 32 |
| UK [89] | WGS | Entire gut | Untreated | Apiary | Untreated | 23 |

<sup>A</sup>We exclusively focused on subjects that included all possible gut sites where bacteria can reside. <sup>B</sup> Using sucrose as a control. <sup>C</sup> Using untreated and sucrose-treated samples as controls.

To address potential batch effects in 16S datasets due to sequencing method heterogeneity [34,35], we performed Principal Coordinates Analysis (PCoA) based on Bray-Curtis dissimilarity. Clustering was primarily driven by variations in amplicon regions (PERMANOVA: R² = 0.135, *p* < 0.001) and library sizes (R² = 0.149, *p* < 0.001) (Fig. S3A). These effects were minimized after trimming libraries to a common V4 region and standardizing library sizes via rarefaction and filtering based on prevalence and abundance thresholds (see Fig. S3B and Methods). Although beta diversity differences related to study comparisons and experiment types (adult vs. newly emerged bees) persisted, we conclude that batch effects from technical variations, especially amplicon region and library size, are negligible after trimming and normalization and can be disregarded when comparing gut microbiome compositions across geographies.

### Honeybee gut microbiota is composed of a stable core with region-depend abundance patterns

We first examined the most abundant taxa in untreated (field control) honeybee gut microbiomes using 16S datasets from Armenia, Denmark, New Zealand, and the southern US, along with WGS datasets from China and the UK. Consistent with prior research, the ten most abundant genera across all samples (both 16S and WGS) included: Bacillota*: Lactobacillus (*formerly *Lactobacillus Firm-5) and Bombilactobacillus (*formerly *Firm-4)*; Pseudomonadota*: Gilliamella, Snodgrassella, Frischella, Bartonella, Bombella, Commensalibacter,* and *Klebsiella;* Actinobacteriota*: Bifidobacterium* [51,52]. Although *Klebsiella,* an opportunistic pathogen, was infrequently detected, it reached an average abundance of 3.2% in some samples (Fig. 2A).

**Figure 2.**
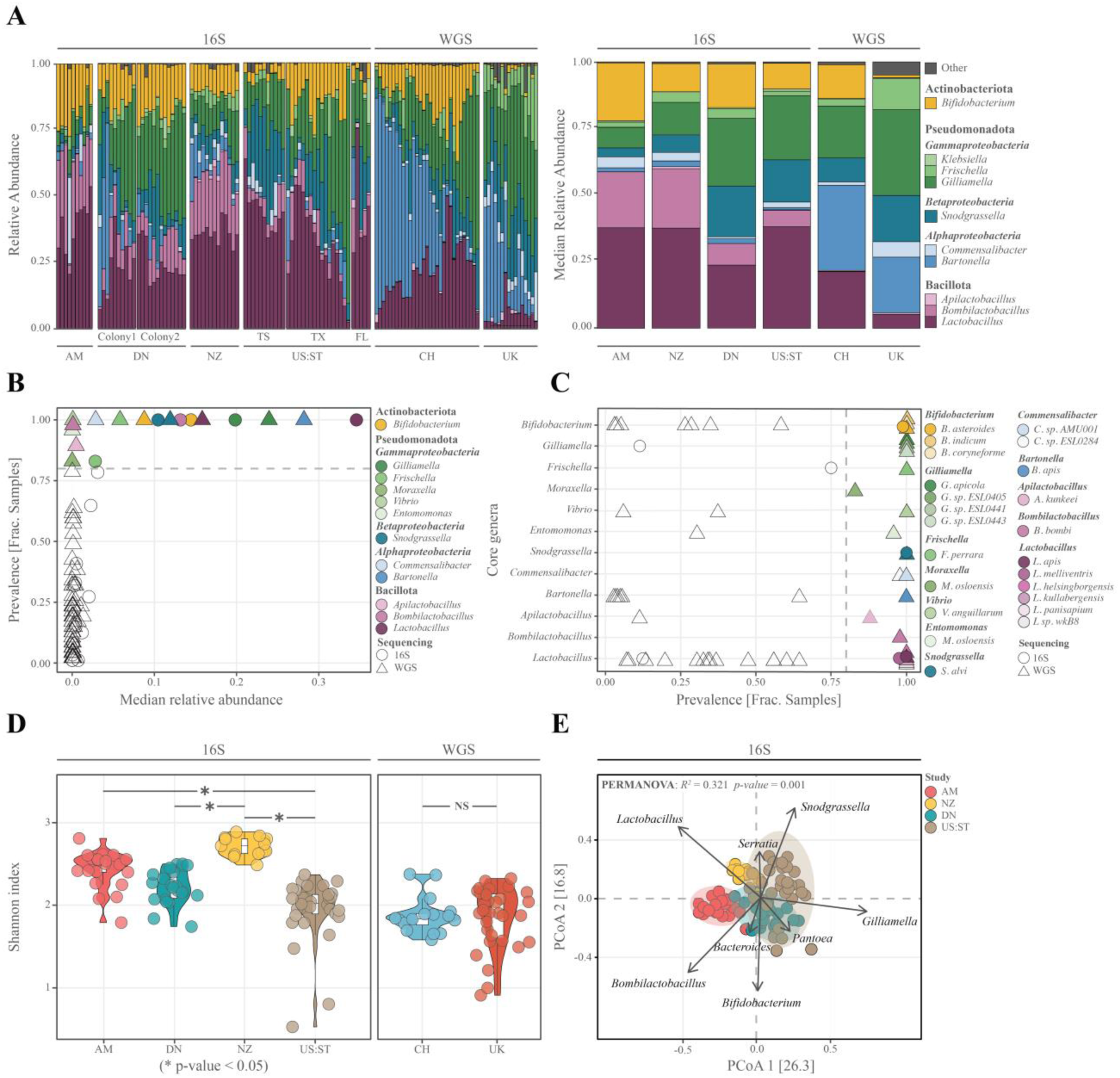
Comparison of honeybee gut microbiome composition across studies. **(A)** Relative abundances of the 10 most abundant bacterial genera in field control samples, with median relative abundances indicated for each study (AM: Armenia, DN: Denmark, NZ: New Zealand, US:ST: Southern US, CH: China, UK: United Kingdom). (**B)** Prevalence and median relative abundance of core genera (present in >80% of field control samples) across studies. **(C)** Prevalence of species-resolved core taxa across all datasets. (**D)** Alpha diversity of the honeybee gut microbiome in field control samples, measured using Shannon index; asterisks indicate significant pairwise differences (Kruskal-Wallis post-hoc test, *p* < 0.05). The central line indicates the median, the box represents the interquartile range (25th–75th percentiles), and whiskers extend to the 10th and 90th. **(E)** Principal coordinate analysis (PCoA) based on Bray-Curtis distances, with samples colored by study; between-group differences were assessed using PERMANOVA. Arrows denote the genera discriminating community composition (multiple regression test with permutations, *p* < 0.05).

The microbiome composition was relatively stable within and between colonies (ANOSIM, R = 0.183, *p* = 0.024), yet varied significantly between states within studies (ANOSIM, R = 0.246, *p* = 0.002) and among studies (ANOSIM, R = 0.366, *p* = 0.001). In 16S datasets, Denmark and the southern US exhibited higher abundances of *Gilliamella* (26% and 24% median relative abundance) and *Snodgrassella* (19% and 16%), while *Bombilactobacillus* (21% and 22%) dominated in Armenia and New Zealand. Armenia also showed elevated *Bifidobacterium* levels. In contrast, WGS datasets were dominated by *Bartonella* (17%-31%), whereas 16S datasets were dominated by *Lactobacillus* (median 22%-42%). *Bartonella* was less prevalent in 16S datasets (0.4%–9%) and absent in the southern US (Fig. 2B).

Next, we defined core taxa as those present in at least 80% of field control samples per study, analyzing 16S and WGS datasets separately (Fig. 2B, C). In the 16S datasets, six core taxa were identified: *Lactobacillus Bombilactobacillus, Gillimella*, *Snodgrassella, Frischella,* and *Bifidobacterium*. WGS datasets revealed six additional core genera: *Entomomonas, Moraxella, Vibrio* (Gammaproteobacteria)*; Bartonella, Apilactobacillus* and *Commensalibacter* (Alphaproteobacteria).

Notably, five core genera, *Lactobacillus, Bombilactobacillus, Gillimella*, *Snodgrassella,* and *Bifidobacterium,* are well documented [11,53,54]. In contrast, *Bartonella, Commensalibacter, Frischella,* and *Apilactobacillus* were reported as core only in specific studies [2,19,55]. In WGS datasets from the UK and China, we also found three low-abundance genera not previously reported as core: *Etomomonas,* a novel genus from Asian honeybee gut [56], *Moraxella,* previously identified in *Apis mellifera* [57] and linked to nicotinoid exposure [24], and *Vibrio,* enriched in pollen [58] (Table S3).

At the species level, in the 16S datasets, *Lactobacillus* resolved to *L. apis* and *L. melliventris*, whereas WGS datasets revealed a broader range of core species, including *L. apis, L. kullabergensis*, *L. helsingborgensis,* and *L. sp. wkB8*. Notably, *L. melliventris* was not identified by Kraken 2 in the China and UK WGS datasets despite its presence among reads, suggesting database annotation issues (see Supplementary Note 1, Fig. S4). Core *Gilliamella* species were resolved only in WGS samples, dominated by *G. apicola* along with *G. sp. ESL0441, ESL0405,* and *ESL0443*. In *Bifidobacterium, B. asteroides* was core in both 16S and WGS, with *B. coryneforme* and *B. indicum* additionally core in WGS. In *Snodgrassella, S. alvi* was the only core species consistently identified in 16S and WGS. Additional WGS-only core species were *Bombilactobacillus bombi, Frischella perrara, Bartonella apis*, *Commensalibacter sp. ESL0284 and AMU001, Apilactobacillus kunkeei, Entomomonas moraniae*, *Moraxella osloensis,* and *Vibrio anguillarum* (Fig. 2C).

We then compared microbiome diversity across studies. Alpha-diversity (Shannon index) was significantly higher in New Zealand compared to Southern US and Denmark (Kruskal-Wallis, post-hoc *p* < 0.001). Armenia also exhibited higher diversity than the Southern US (*p* < 0.001) (Fig. 2D, Table S4). PCoA using Bray-Curtis dissimilarity revealed that New Zealand and Armenia samples clustered distinctly from other studies, indicating greater within-group homogeneity (Fig. 2E, Table S5).

The features influencing discrimination of Armenia and New Zealand samples were increased *Lactobacillus* and *Bombilactobacillus*, and decreased *Gilliamella* (Fig. 2E). Notably, a subgroup of Southern US samples from Texas differed from those in Florida and Tennessee due to higher *Pantoea* abundance (*e.g. P. sp. SO10, P. vagans, P. agglomerans, P. ananatis*). These species may act as plant pathogens or contribute to pathogen defense in both plants and bees, with some linked to urban areas [59–63].

Overall, our analysis reveals a conserved core microbiota across diverse geographic regions, although their relative abundances vary significantly by location. We next investigated how seasonal and environmental factors shape the gut microbiota of honeybees in Armenia.

### Seasonal and environmental factors influence gut microbiota in Armenia, with increased *Commensalibacter* abundance in autumn

We next analyzed original data from Armenia, where the gut microbiomes of worker honeybees were collected from nine regions during 2017, 2020, and 2023, in spring, summer, and autumn. In total, 21 samples, each comprising 5-6 pooled bee guts were examined.

The microbiome was largely dominated by *Lacotibacillus, Bombilactobacillus*, and *Bifidobacterium* (Fig. 3A). Notably, some samples exhibited increased levels of potential pathogens such as *Escherichia*, and *Shigella*.

**Figure 3.**
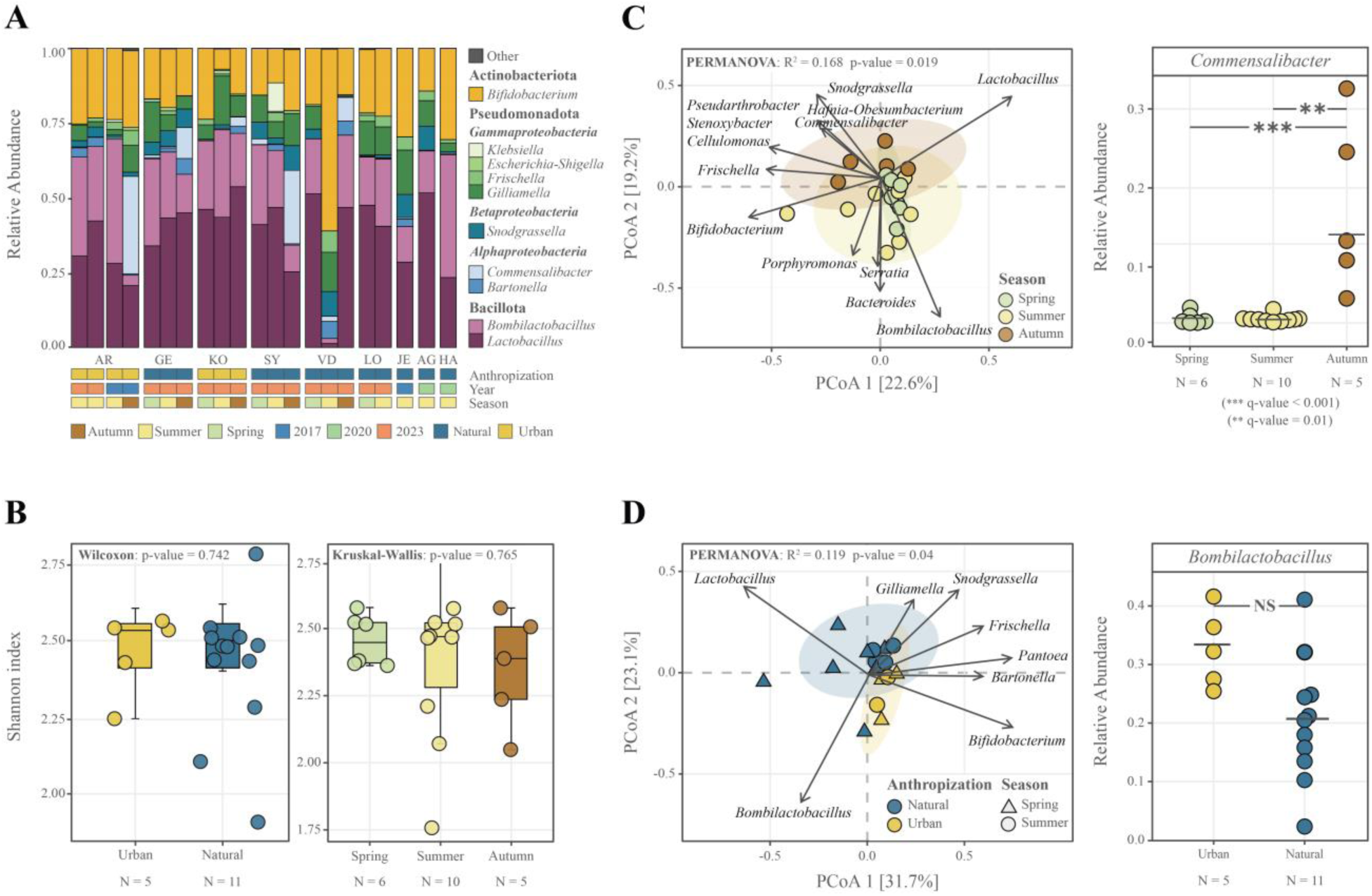
Impacts of seasonality and urbanization on honeybee gut microbiome composition in Armenia. **(A)** Relative abundance of major taxa in Armenian honeybee gut samples, stratified by season, collection date, and anthropization level. **(B)** Alpha diversity of the gut microbiome (Shannon index) analyzed according to seasonal and anthropization factors. The central line indicates the median, the box represents the interquartile range (25th– 75th percentiles), and whiskers extend to the 10th and 90th. **(C)** Principal coordinate analysis (PCoA) based on Bray–Curtis distances; arrows indicate the most discriminant genera (multiple regression with permutations, *p* < 0.05). Samples are colored by collection season. The adjacent scatterplot highlights seasonal differences in *Commensalibacter* abundance, with group differences determined using ANCOM-BC2. **(D)** PCoA is colored by anthropization level, with an accompanying scatterplot illustrating differences in *Bombilactobacillus* abundance between urban and natural habitats.

Seasonal comparisons among spring, summer, and autumn samples revealed significant differences between autumn and summer (Fig 3C; PERMANOVA, R^2^ = 0.155, *p* = 0.03, Table S6). Multiple regression analysis indicated that these differences were primarily driven by higher abundances of *Commensalibacter* and *Snodgrassella* in autumn, and increased *Bombilactobacillus* in summer. Differential abundance analysis using ANCOM-BC2 confirmed a statistically significant rise in *Commensalibacter* during autumn (summer-autumn, *q* < 0.001, LFC = 3.94; spring-autumn, *q* < 0.016, LFC = 4.56) (Fig. 3C, Table S7). Similar seasonal shifts have been observed elsewhere, with Swiss studies reporting increased *Commensalibacter* in winter bees [64], although other European [65] and Chinese studies have noted alternative patterns, e.g. increase in *Bartonella* [66] or a decrease in *Snodgrasella* [67].

We further assessed anthropogenic impacts by comparing bees from urban/near-urban apiaries (Ararat and Kotayk, near Yerevan) with those from rural or natural regions. PCoA revealed distinct microbiota profiles between urban and natural sites (Fig. 3D; PERMANOVA, R² = 0.119, p = 0.04), primarily due to an increased abundance of *Bombilactobacillus* in the urban samples. However, differential abundance analysis with ANCOM-BC2 did not yield statistically significant differences (q < 1, LFC = 0.32), likely due to the small sample size (Fig. 3D, Table S7). To date, no specific taxon has been consistently linked to anthropogenic factors in urban regions. However, a decrease of *Lactobacillus* and *Snodgrasella,* with a simultaneous increase in *Bifidobacterium* and *Enterobacteriacea,* seen in near-urban regions, although not significant, was also reported previously [68]. Although some studies have reported increased microbial diversity in urban environments [61,69,70], we did not observe such an effect (Fig. 3B).

### Differential Impact of Pesticides and Diet on Honeybee Gut Microbiome

We investigated the impact of pesticides, herbicides, and dietary supplements on the diversity and composition of the honeybee gut microbiome. To ensure comparability, all datasets were processed using a uniform preprocessing and analysis pipeline. In the studies considered, treatments were administered via feeding, with controls provided either as sugar water (sugar controls [19]) or as field controls, depending on the study. Comparisons were made between treated samples and their corresponding controls.

Oxalic acid treatment produced the most pronounced shift in beta diversity (PERMANOVA, R² = 0.197, *p* = 0.001), while alpha diversity did not differ significantly from sugar controls (Fig. 4A). Taxonomic analysis using ANCOM-BC2 revealed that oxalic acid was associated with significant decreases in *Bombella intestini* (logFC < -3.8, adjusted *p* < 0.001)*, Bartonella* (logFC < -1.4, adjusted *p* = 0.003)*, Apilactobacillus* (logFC < -1.4, adjusted *p* = 0.003), alongside an increase in *Klebsiella* (logFC > 1.2, adjusted *p* = 0.013) (Fig. 4E, Table S8). These findings are consistent with the original study [19], which employed the ALDEx2 method [71] and detected similar taxa, although additional species were reported.

**Figure 4.**
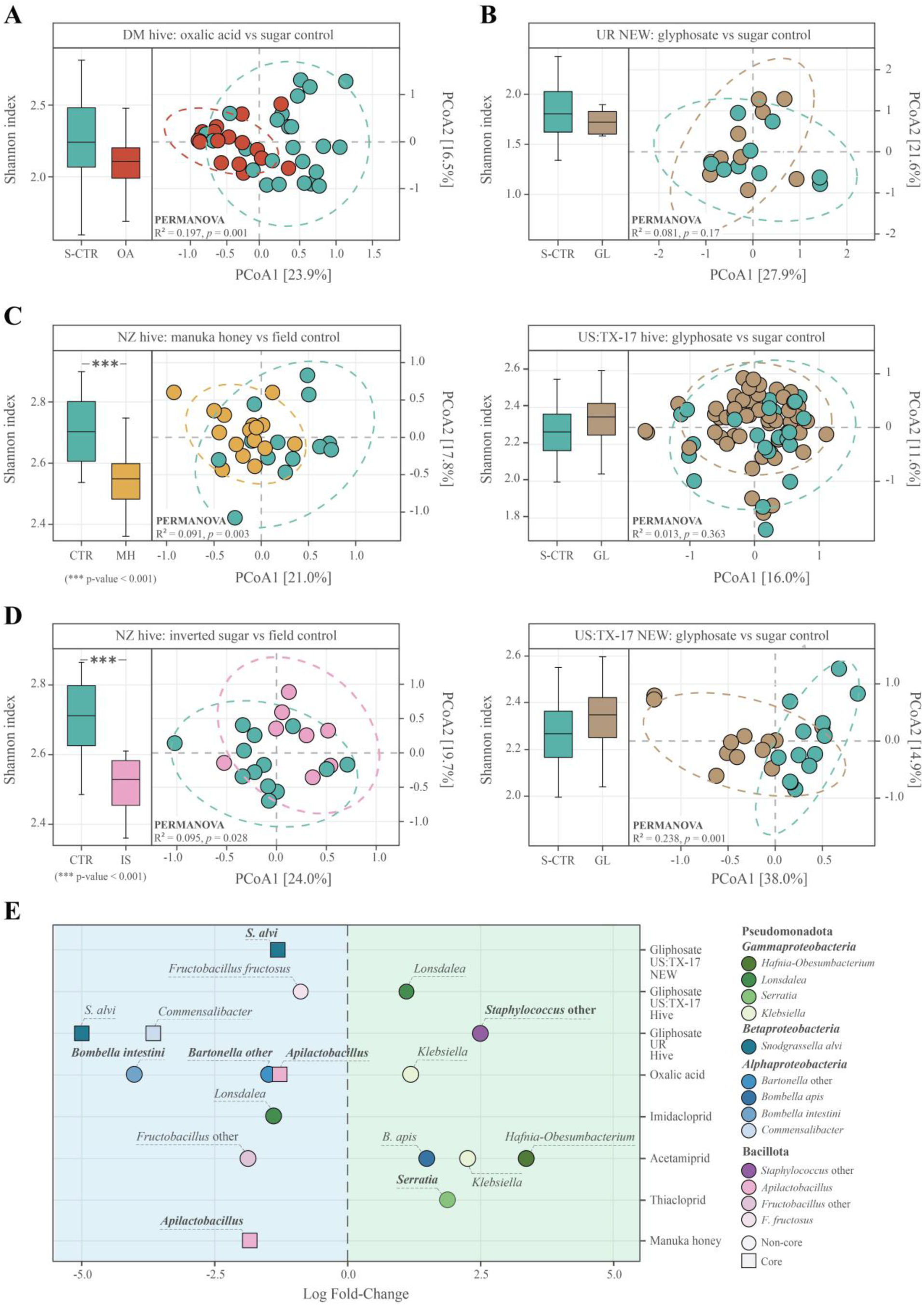
Effects of treatments on alpha diversity, community composition, and differential abundance of specific taxa. **(A–D)** Changes in the honeybee gut microbiome following various treatments. Alpha diversity (Shannon index) is presented as boxplots, where the central line indicates the median, the box represents the interquartile range, and whiskers extend to the 10th and 90th percentiles. Group differences in alpha diversity were assessed using the Kruskal-Wallis test (p-values indicated). Beta diversity was evaluated using Bray-Curtis distances and visualized via PCoA, with ellipses representing 95% confidence intervals; PERMANOVA (R² and p-values) was used to assess differences between groups. The panels depict: **(A)** Oxalic acid (vs. sugar controls); **(B)** Glyphosate treatment in adult honeybees from Uruguay and Texas, USA, as well as newly emerged bees in Texas (vs. sugar controls); **(C)** Manuka honey (vs. field controls); and **(D)** Inverted sugar (vs. field controls). **(E)** Differentially abundant taxa identified by ANCOM-BC2 following treatment. Log₂ fold changes (logFC) are plotted on the x-axis, and only taxa with statistically significant differences (adjusted p-value < 0.05) are shown. Taxa names in bold indicate those that passed the sensitivity analysis for pseudo-count addition. Squared taxa belong to the core, round taxa are non-core.

In contrast, treatments with other insecticides, including acetamiprid, thiacloprid, and imidacloprid did not lead to significant shifts in overall microbiome composition (Fig. S6), consistent with previous studies [4,19,25]. Nonetheless, differential abundance analysis identified specific ASVs that were altered by these insecticides, including increased abundances of *Bombella apis, Hafnia-Obesumbacterium, Klebsiella*, *Lonsdalea,* and *Serratia*, and a decrease in *Fructobacillus* and *Apilactobacilluse* (Fig. 4E, Table S8). We did not detect significant composition differences for other insecticides evaluated in the same study [19] (Fig. S6).

For glyphosate, we analyzed data from studies conducted in Uruguay [72] and Texas, USA [29]. In the Texas study, two experimental setups were performed, one treating adult bees and another treating newly emerged bees (NEW samples), while in the Uruguay study, only NEW samples were considered. Beta diversity analysis revealed a significant shift in Texas NEW samples (PERMANOVA, R^2^ = 0.238, *p* = 0.001). In contrast, no significant changes were detected in adult bees from Texas or NEWs from Uruguay, in agreement with the original reports [4,29]. Although the Uruguay study reported compositional changes in NEW bees, these findings were based on comparisons involving multiple treatments rather than glyphosate alone (Table S9). Notably, ANCOM-BC2 analysis showed consistent reductions in *Snodgrasella alvi* in both the Texas and the Uruguay NEW samples (Fig. 4E, Table S8) [4,29]. In adult bees from Texas, glyphosate treatment also resulted in a decrease in *Fructobacillus fructosus* and an increase in *Lonsdalea*, whereas the Uruguay NEW revealed reductions in *Commensalibacter* and *S. alvi* coupled with an increase in *Staphylococcus*.

In contrast to pesticide treatments, dietary supplements such as manuka honey and inverted sugar significantly decreased alpha diversity relative to field controls, consistent with previous findings from New Zealand [18]. Both supplements induced shifts in microbiome composition as determined by beta diversity analysis, though the effect sizes were smaller than those observed with oxalic acid and glyphosate in Texas NEW samples (PERMANOVA R^2^ = 0.091, *p* = 0.003 for manuka honey; R^2^ = 0.095, *p* = 0.003 for inverted sugar) (Fig. 4C, D). At the taxonomic level, manuka honey supplementation was associated with a decrease in *Apilactobacillus*, a finding not reported in the original study (Fig. 4E, Table S8, S9). Additionally, we did not identify any differentially abundant taxa with inverted sugar treatment, in contrast to the original study, which utilized ANOVA for statistical analysis [18].

Collectively, these results highlight that different treatments distinctly modulate the honeybee gut microbiome, with some (e.g., oxalic acid) eliciting robust compositional shifts and others (e.g., certain insecticides) causing more subtle, taxon-specific changes. However, given that focusing solely on individual taxa may not fully capture the complexity of the microbial community’s adaptations, we next employ co-abundance network analysis to elucidate collective effects and functional responses to these treatments.

### Co-Abundance Networks Reveal Adaptive Pesticide Responses and Potential Glyphosate Degradation in Honeybee Gut Microbiome

To capture community-level responses to pesticide and sugar supplement treatments, we used the microbiome abundance data of all the studies and constructed a co-abundance network where each taxon is represented as a node and edges denote significant positive or negative correlations in their relative abundances. We then identified the subnetwork in which the sum of log fold change differential abundance weights (compared to controls) was greatest, thereby pinpointing the group of taxa that collectively respond most strongly to the treatment. Finally, we performed functional enrichment analysis on this subnetwork to determine which pathways and functional profiles may enable these bacteria to adapt and increase in abundance (Fig. 5 and Fig. S6; see Methods for details).

**Figure 5.**
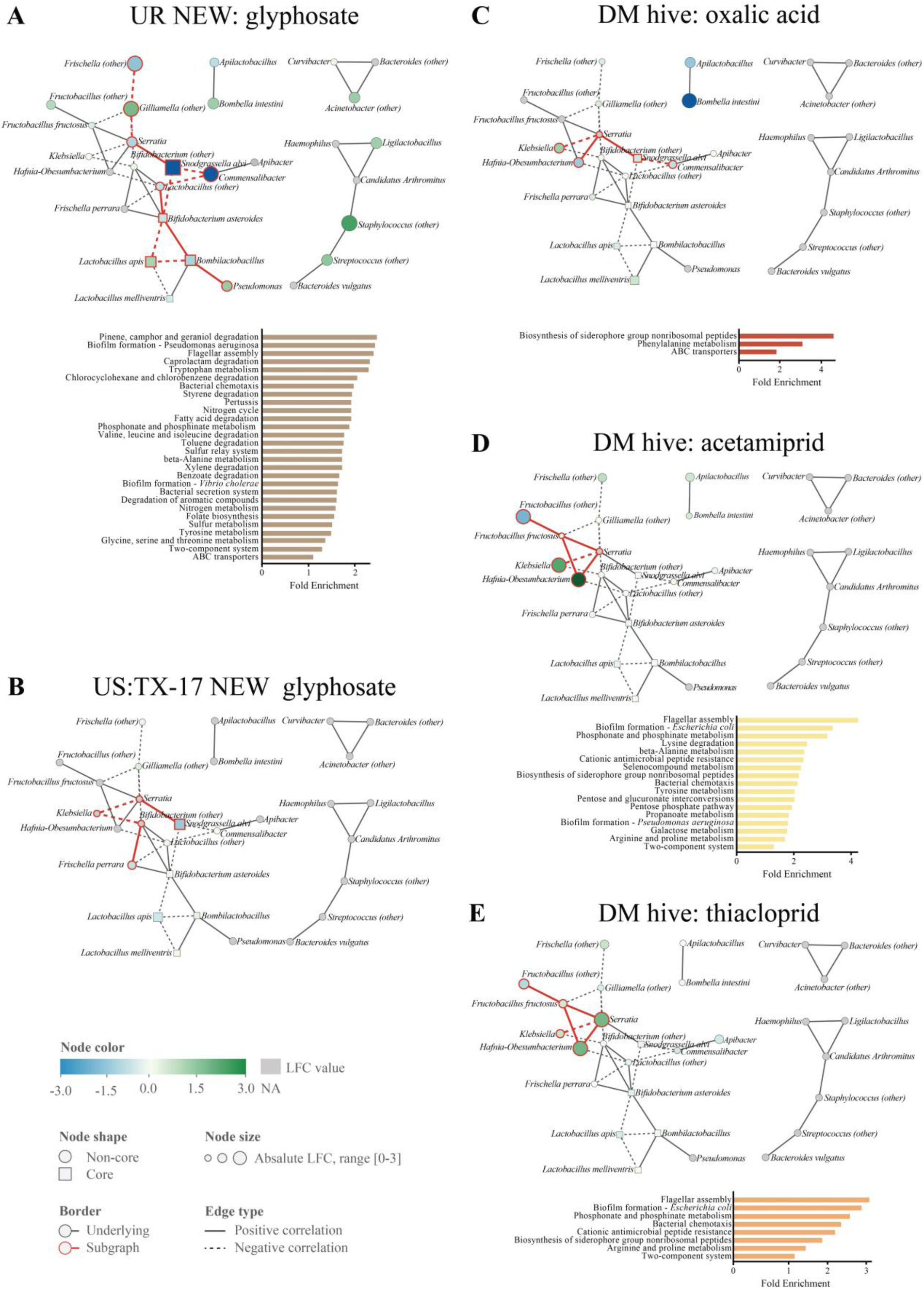
Co-abundance networks of the honeybee gut microbiome under treatments and corresponding KEGG pathway overrepresentation analysis. Panels depict treatment-associated subnetworks for: **(A)** glyphosate in Uruguay hive samples, **(B)** glyphosate in US Texas NEW samples, **(C)** oxalic acid, **(D)** acetamiprid, and **(E)** thiacloprid. For each treatment, the upper panel shows the co-abundance network and the lower panel displays the corresponding functional enrichment results. In the networks, nodes represent taxa and edges denote significant Spearman correlations (ρ > 0.6, adjusted p < 0.2); solid lines indicate positive correlations, and dashed lines negative correlations. Core taxa are represented by rectangular nodes, and non-core taxa by circles. Node size reflects the absolute log fold change (LFC) between treatment and control groups, with a green-to-blue color scale indicating up- and down-regulation, respectively. Nodes for which LFC values were not calculated by ANCOM-BC2 due to sample size limitations are colored gray. Nodes and edges highlighted with red borders indicate the treatment-specific subnetwork identified using the maximum weight connected subgraph algorithm (see Methods). Only KEGG pathways significantly enriched in the treatment groups (FDR < 0.1) are shown.

The glyphosate-associated subgraph identified in the Uruguay and Texas NEW samples included interconnected taxa, with decreases in *S. alvi* in Uruguay and Texas, and decrease in *Serratia* in Uruguay. In the Uruguay NEW samples, this subnetwork expanded to encompass additional taxa: *Frischella*, *Commensalibacter*, *Lactobacillus*, *Bifidobacterium asteroides*, and *Bombilactobacillus* (all decreased), along with increases in *Gilliamella*, *Fructobacillus* (other), *Lactobacillus apis* and *Pseudomonas*.

Pathway enrichment within this glyphosate-associated subnetwork in Uruguay identified several amino acid metabolic pathways, such as tryptophan and tyrosine metabolism, suggesting a direct influence of glyphosate exposure (Fig. 5A, B, Table S12). Furthermore, our analysis revealed an enrichment of pathways for xenobiotic and aromatic compound degradation, nitrogen cycling, and two-component system, suggesting potential capacity for glyphosate degradation. *Pseudomonas* is known to degrade glyphosate either by converting it to sarcosine via the C-P lyase pathway or by transforming it into aminomethylphosphonic acid (AMPA) and methylamine [73–76]. In our samples, we observed enrichment of the phosphonate and phosphinate metabolism pathway, including the *phnJ* gene, which encodes C-P lyase; a key enzyme in both degradation routes. This gene is part of the phn operon, which also encodes putative glyphosate transporters (*phnC*, *phnD*, and *phnE*) [77], indicating glyphosate uptake capacity. Additionally, enriched pathways for glycine metabolism, nitrogen cycling, and aromatic compound degradation may further facilitate the breakdown of glyphosate degradation intermediates. Although the precise degradation route remains unclear, the increased abundance of *Pseudomonas* and these functional enrichments support the hypothesis that glyphosate degradation is an adaptive response to glyphosate exposure. In Uruguay samples, glyphosate exposure altered bacterial chemotaxis and biofilm formation pathways, likely mediated by both *Pseudomonas* and *Gilliamella*. This response may enhance overall bacterial stress adaptation [10]. Furthermore, a disconnected subnetwork in these samples showed upregulation of *Staphylococcus* and *Streptococcus*, aligning with previous reports of impaired immune function following glyphosate treatment [78].

In contrast, treatments with oxalic acid and neonicotinoids (acetamiprid and thiacloprid) affected a different subnetwork that included *Klebsiella*, *Hafnia-Obesumbacterium*, and *Serratia*. *Klebsiella* and *Serratia* increased in both cases, while *Hafnia-Obesumbacterium* increased under neonicotinoid exposure, and decreased under oxalic acid treatment. (Fig. 5C). For neonicotinoids, this subnetwork further expanded to include *Fructobacillus*, which decreased in abundance (Fig. 5D,E). Oxalic acid’s impact also extended to include decreased abundances of *Snodgrassella alvi,* and *Commensalibacter*, highlighting the documented sensitivity of these taxa to pesticides [19].

Unlike glyphosate, we did not detect specific oxalic acid degradation pathways in the samples. Instead, the oxalic acid–affected subnetwork showed enrichment for stress response pathways. ABC transporters could enhance the regulation of internal pH after its reduction by oxalic acid application [79]. Upregulation of siderophore biosynthesis suggests a compensatory response to iron limitation caused by oxalic acid sequestration [80]. Similarly, neonicotinoid exposure commonly enriched pathways related to stress responses, such as the two-component system, which is typically involved in insecticide and xenobiotic detoxification [81,82]. Notably, the pathway for cationic antimicrobial peptide (CAMP) resistance was enriched in neonicotinoid-associated subnetworks, potentially reflecting increased antimicrobial peptide production by the host, as previously observed in bumblebees exposed to these insecticides [72] (Fig. 5D-E, Table S12).

## Discussion

A major strength of our study is the standardized re-analysis of diverse datasets. By using both 16S and WGS data and a unified pipeline, we minimize methodological discrepancies that may have contributed to conflicting findings in previous studies [2,34,35]. Our novel co-abundance network approach links compositional shifts with functional potential. We also provide a fully reproducible pipeline to facilitate future research across diverse biomes of interest, along with an R package, *micronet,* for co-abundance network analysis.

Nevertheless, we acknowledge several limitations. Despite efforts to normalize library size and limit primer biases, residual heterogeneity remained across studies, especially due to differing extraction methods, primers, and sequencing approaches. This, however, is an inherent issue of any meta-analysis. Additionally, variations in laboratory and field pesticide exposures, cross-sectional sampling designs, and the inherent limitations of ASV-based annotations (which miss strain-level differences) constrain our interpretations. Notably, the WGS data were available only for untreated samples, limiting strain-level functional inference in treated groups.

Our results support the consensus that honeybees maintain a defined core gut microbiome, typically including six phylotypes: *Lactobacillus, Bombilactobacillus, Gilliamella, Snodgrassella, Frischella*, and *Bifidobacterium*. Notably, incorporating low-abundance taxa via WGS datasets suggests an expanded core of up to twelve taxa, including *Bartonella, Commensalibacter, Frischella, Apilactobacillus, Moraxella,* and *Vibrio*.

Studies show that honeybee microbiota are shaped by diet, climate, and local flora [67,68,83–85]. We also observed significant geographic variation. However, the factors driving this variation are challenging to disentangle. Seasonal factors and urbanization likely play a role, as evidenced by our newly generated data from Armenia. Other biological factors unaccounted for in our study, such as bee age, temperature, floral resources, and colony management, may also contribute to these differences. This may explain why different studies often conflict; for instance, our finding of elevated *Commensalibacter* in autumn agrees with a Swiss study but contrasts with reports from other European and Asian sites [64,66]. Moreover, variations in data analysis pipelines have contributed to inconsistent results, which underscores the value of our unified reprocessing of both 16S and WGS datasets from multiple studies.

A pressing question in bee research is how agrochemicals affect colony health. Recent studies link insecticides, and herbicides to shifts in gut community composition, with conflicting or inconclusive findings regarding affected taxa and functions [4,19,23,25]. Our meta-analysis supports that gut responses vary by chemical class, dose, and bee life stage.

Glyphosate affected the core taxa. We consistently observed depletion of *Snodgrassella alvi* confirming its reported sensitivity. In Texas, newly emerged bees showed marked beta diversity shifts, and in Uruguayan newly emerged bees exhibited drastic network changes suggesting higher vulnerability in younger bees. Adult bees in Texas displayed milder shifts, hinting at a more resilient microbiome in older foragers [4,29].

Co-abundance network analysis revealed potential adaptation mechanisms. In Uruguayan newly emerged bees, a glyphosate-affected subnetwork involving increased *Pseudomonas, Gilliamella,* and enrichment of several pathways, suggesting a potential for C-P lyase-mediated glyphosate degradation. While *Pseudomonas* species have demonstrated glyphosate utilization in other environments [76,77], this, to the best of our knowledge, is the first indication of such an adaptation mechanism in honeybees. Enrichment of tryptophan and tyrosine metabolism pathways further suggests compensation for glyphosate-induced shikimate pathway inhibition.

In contrast, the Texas newly emerged bees exhibited a smaller affected subnetwork. Strain-level differences or variations in experimental setup could affect these differences. The Texas bees were exposed to a higher glyphosate dose (169 mg/l versus 10 mg/l in Uruguay), however at a shorter period (2 days versus 7 days in Uruguay). This also suggests that prolonged exposures have a stronger impact on microbiome shifts and adaptations than higher doses.

Insecticides, on the other hand, did not affect the core taxa but induced an increased abundance of opportunistic pathogens that exist as part of the normal microbiota. Neonicotinoids (imidacloprid, thiacloprid, acetamiprid) induced subtler microbiome alterations [19]. Although overall community composition remained relatively stable, low-abundance opportunistic pathogens, such as *Serratia*, *Klebsiella*, and *Hafnia-Obesumbacterium* thrived under treatment. These taxa were enriched for pathways of xenobiotic detoxification and stress response (e.g., two-component system, ABC transporters), and cationic antimicrobial peptide (CAMP) resistance. These adaptations could confer a selective advantage if the host produces more antimicrobial peptides under pesticide-induced stress [72].

In contrast, the acaricide oxalic acid caused the most pronounced beta-diversity shift, and also increased *Serratia*, and *Klebsiella*, similar to neonicotinoids. Additionally, it led to the depletion of *Bartonella, Bombella, Apilactobacillus*, and *Snodgrassella* [19,80].

Although considered relatively “bee-safe,” oxalic acid lowers gut pH, imposing selective pressure that appears to upregulate general pathways of response to low pH and iron limitation (e.g., ABC transporters, siderophore biosynthesis), rather than dedicated oxalate-degradation mechanisms [80,86].

All in all, our meta-analysis reaffirms a stable honeybee core microbiota while demonstrating that environmental factors (geography, season) and anthropogenic influences (pesticides, dietary additives) induce taxon-specific shifts. Pesticides tend to favor bacteria with enhanced stress-response capabilities or opportunistic degraders. Longitudinal metagenomics and mechanistic studies remain essential to clarify how these microbiome shifts impact colony health, ultimately guiding sustainable strategies for apiculture and pollinator conservation.

## Data and software availability

Raw 16S rRNA gene sequencing and WGS data sets evaluated in this study are available from the NCBI Sequence Read Archive (PRJNA719169, PRJNA732842, PRJNA775827, PRJNA432211, PRJNA432210, PRJNA483763, PRJNA531038, PRJNA685398, PRJNA787435 and PRJNA494922). The 16s amplicon datasets from the Armenian samples generated in this study are deposited in the NCBI SRA database (PRJNA1220608).

The *micronet* package for co-abundance network analysis is available for download and use at Github: https://github.com/abi-am/micronet. The rest of the scripts used in this study are available at https://github.com/abi-am/bee-microbiome-metaanalysis.

## Acknowledgements

The authors acknowledge funding from the Higher Education and Science Committee (HESC) MESCS RA PhD Support program awarded to NV (24AA-1F065), the HESC RA Prospective Directions grant awarded to LN (24FP-2I061), and Yerevan State University Inner Grant 2022 to IB.

## Contributions

LN and NV conceived the study. NV performed the analyses, produced the figures, and wrote the manuscript. IB generated the Armenian data. LA developed the *micronet* package and conducted the co-abundance network analysis. MZ and HL carried out additional experiments. LN supervised the study and prepared the manuscript. CM supervised computational analysis and revised the manuscript. All authors read and approved the final version of the manuscript.

