## Supplementary material for "Global distribution of honeybee gut microbiome and pesticide-driven adaptations in opportunistic microbial species": Supplementary Figure Legends.docx

**Figure S1.** Rarefaction curves illustrating the number of identified ASVs as a function of library size for each study. Dashed lines represent the rarefaction **(A)** threshold of 12 000 reads for 16S rRNA and **(B)** 800 000 for WGS, used to normalize sequencing depth across samples.

**Figure S2.** Principal coordinate analysis (PCoA) of Bray-Curtis dissimilarity, colored by hypervariable regions of the 16S rRNA gene **(A)** before and **(B**) after trimming the libraries to a common V4 region and applying library size reduction using rarefaction, colored by the amplicon region, library size, study and experiment type.

**Figure S3. (A)** Sankey diagram produced by Pavian R package with the total number of corrected reads as estimated with Kraken 2 for all sites combined at the level of kingdom, phylum, family, genus and species. Mean relative coverage of MAGs within WGS studies for **(B)** *Lactobacillus kullabergensis ESL0186* and **(C)** *Lactobacillus melliventris Hma 8*.

**Figure S4.** Alpha and beta diversity analyses showed that the treatments did not alter the structure and membership of the gut microbiota based on Shannon index and Bray-curtis dissimilarities **(A)** acetamiprid and thiacloprid, **(B)** imidacloprid and **(C)** methylglyoxal and dihydroxyacetone (*p* > 0,05 in all cases, **A-C**).

**Figure S5.** Co-abundance networks of the honeybee gut microbiome under **(A)** Dihydroxyacetone and **(B)** Inverted sugar. For each treatment, the upper panel shows the co-abundance network and the lower panel displays the corresponding functional enrichment results. In the networks, nodes represent taxa and edges denote significant Spearman correlations (ρ > 0.6, adjusted p < 0.2); solid lines indicate positive correlations, and dashed lines negative correlations. Core taxa are represented by rectangular nodes, and non-core taxa by circles. Node size reflects the absolute log fold change (LFC) between treatment and control groups, with a green-to-blue color scale indicating up- and down-regulation, respectively. Nodes for which LFC values were not calculated by ANCOM-BC2 due to sample size limitations are colored gray. Nodes and edges highlighted with red borders indicate the treatment-specific subnetwork identified using the maximum weight connected subgraph algorithm (see Methods).
