## Supplementary material for "Global distribution of honeybee gut microbiome and pesticide-driven adaptations in opportunistic microbial species": Supplementary note.docx

### Supplementary note 1

### *Lactobacillus melliventris* misclassification

The absence of *L. melliventris* in WGS datasets, despite its documented presence in honeybee microbiomes and consistent detection in 16S rRNA datasets, raised questions about potential misclassification in the bioinformatics pipeline. To address this, we conducted a series of targeted analyses to identify the underlying cause. To reject the absence of the L. melliventris genome in the WGS datasets we constructed a custom reference genome database including *L. melliventris* and other representative *Lactobacillus* species from NCBI. Alignment with bowtie2 (v. 2.4.4) confirmed that a lot of reads (48.1% in China and 23.2% in UK) from WGS datasets mapped to the *L. melliventris* *Hma 8* genome, suggesting that its absence in initial classifications was not due to its complete absence in the data (Fig. S3).

Afterwords, to investigate potential classification errors in Kraken2, with the art_illumina tool (v.2016.06.05) [1] we artificially fragmented the reference genomes of *L. melliventris* representative strain *Hma8* and *L. kullabergensis ESL0186* (another highly abundant *Lactobacillus* strain identified in our sample with Kraken 2), creating paired-end FASTQ files with 5x coverage. Kraken analysis of these files,showed that while *L. kullabergensis ESL0186* reads were classified correctly, *L. melliventris Hma8* reads were predominantly misclassified as *Lactobacillus sp. IBH004*, an unclassified *Lactobacillus* species (Fig. S3). However, we did not find *Lactobacillus sp. IBH004* in the WGS samples either.

Further analysis using the kraken2-inspect tool verified that the taxonomic identifiers for both species were accurate and consistent with NCBI Taxonomy. Reference genome quality checks revealed high completeness and low contamination for both *L. melliventris Hma 8* (97.74% completeness, 0.99% contamination) and *Lactobacillus sp. IBH004* (97.68% completeness, 0.81% contamination), ruling out poor assembly or contamination as potential causes.

We conclude that database annotation inconsistencies can significantly impact taxonomic accuracy, as demonstrated by the misclassification of *L. melliventris* in the WGS dataset. The absence of *Lactobacillus sp. IBH004* in the samples further suggests that these misclassifications were due to limitations in the reference database, rather than the biological presence of this species.
