## Supplementary figures and images for "Global distribution of honeybee gut microbiome and pesticide-driven adaptations in opportunistic microbial species"

### Fig.S1.jpg

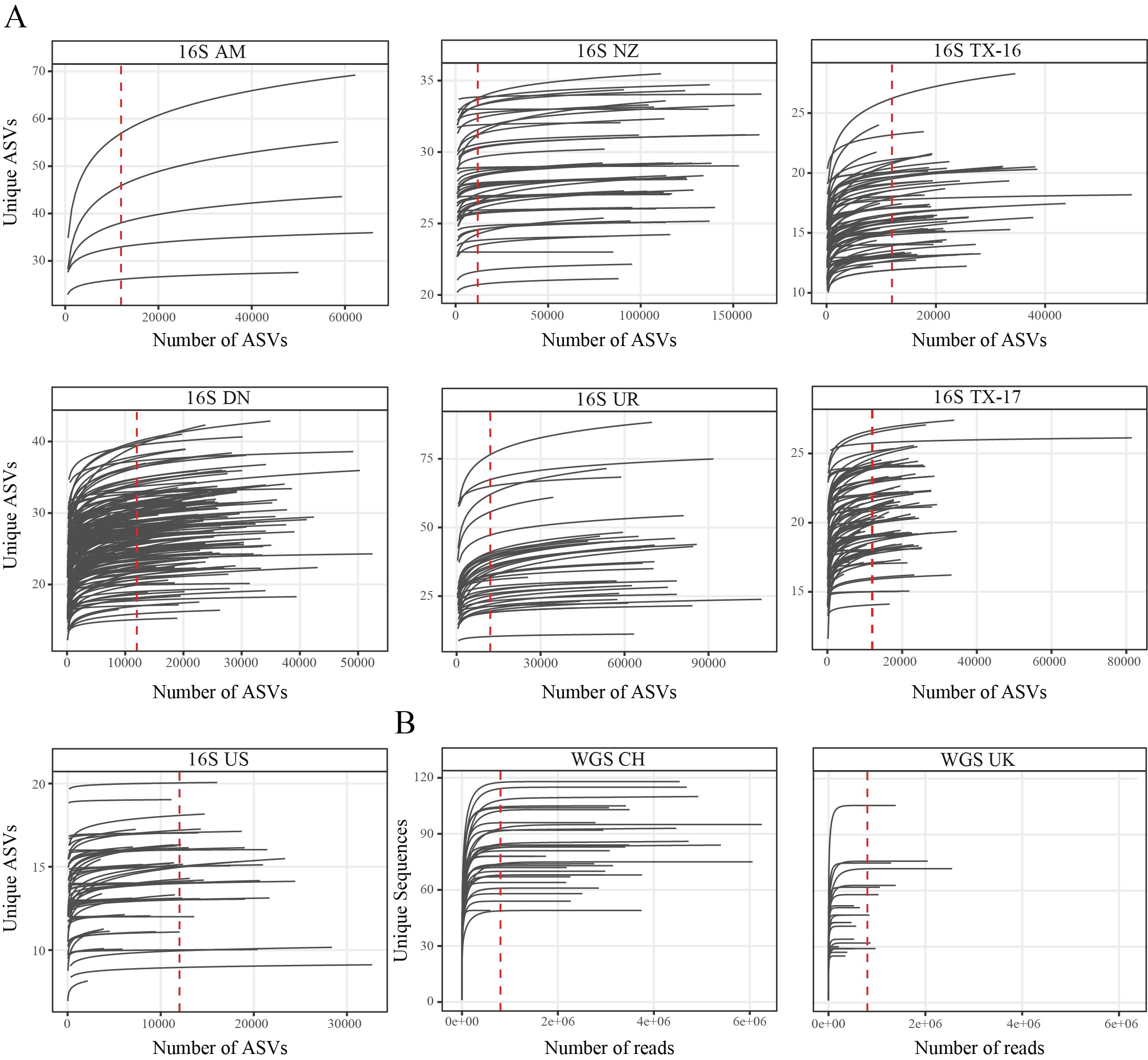

### Fig.S2.jpg

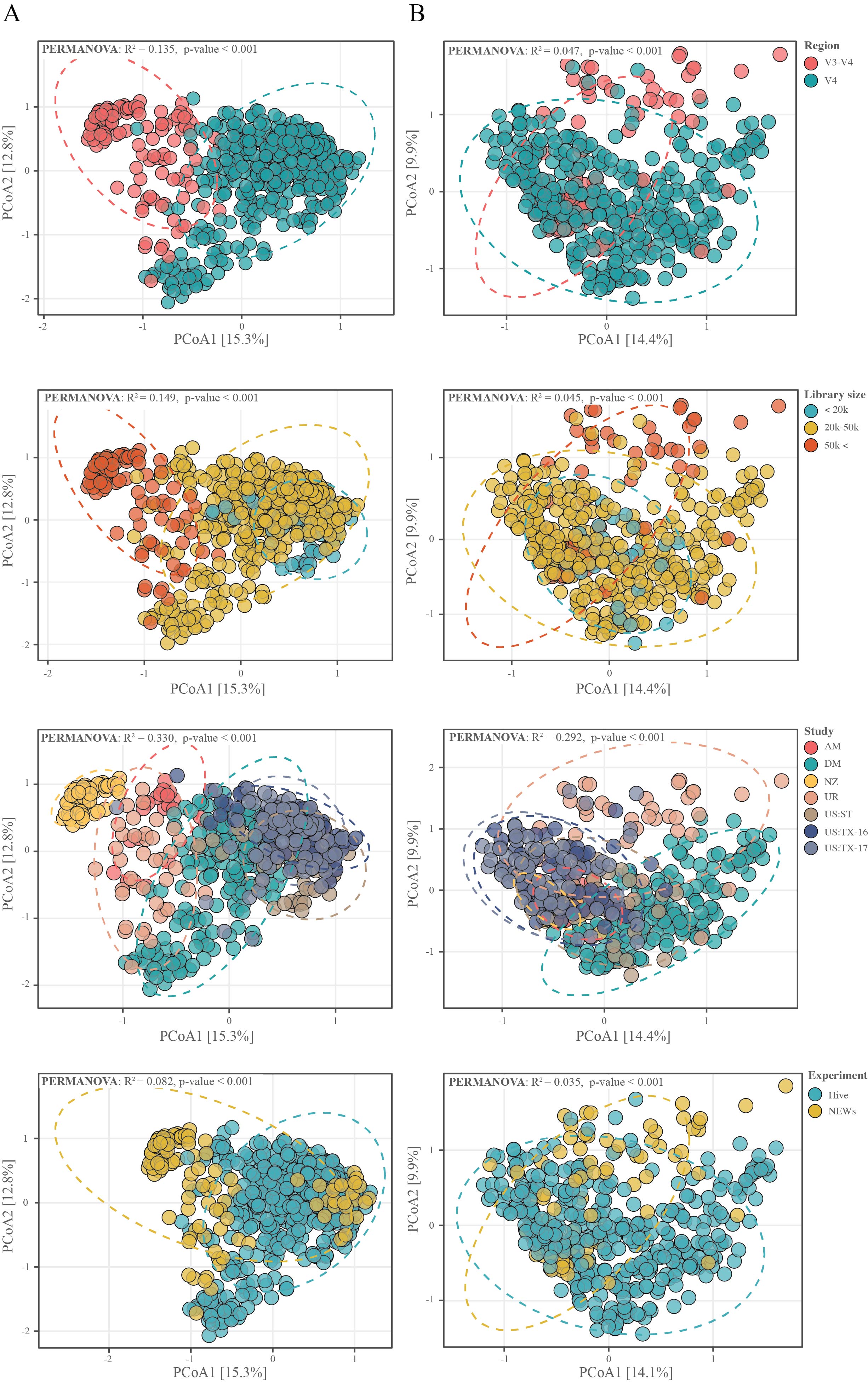

### Fig.S3.jpg

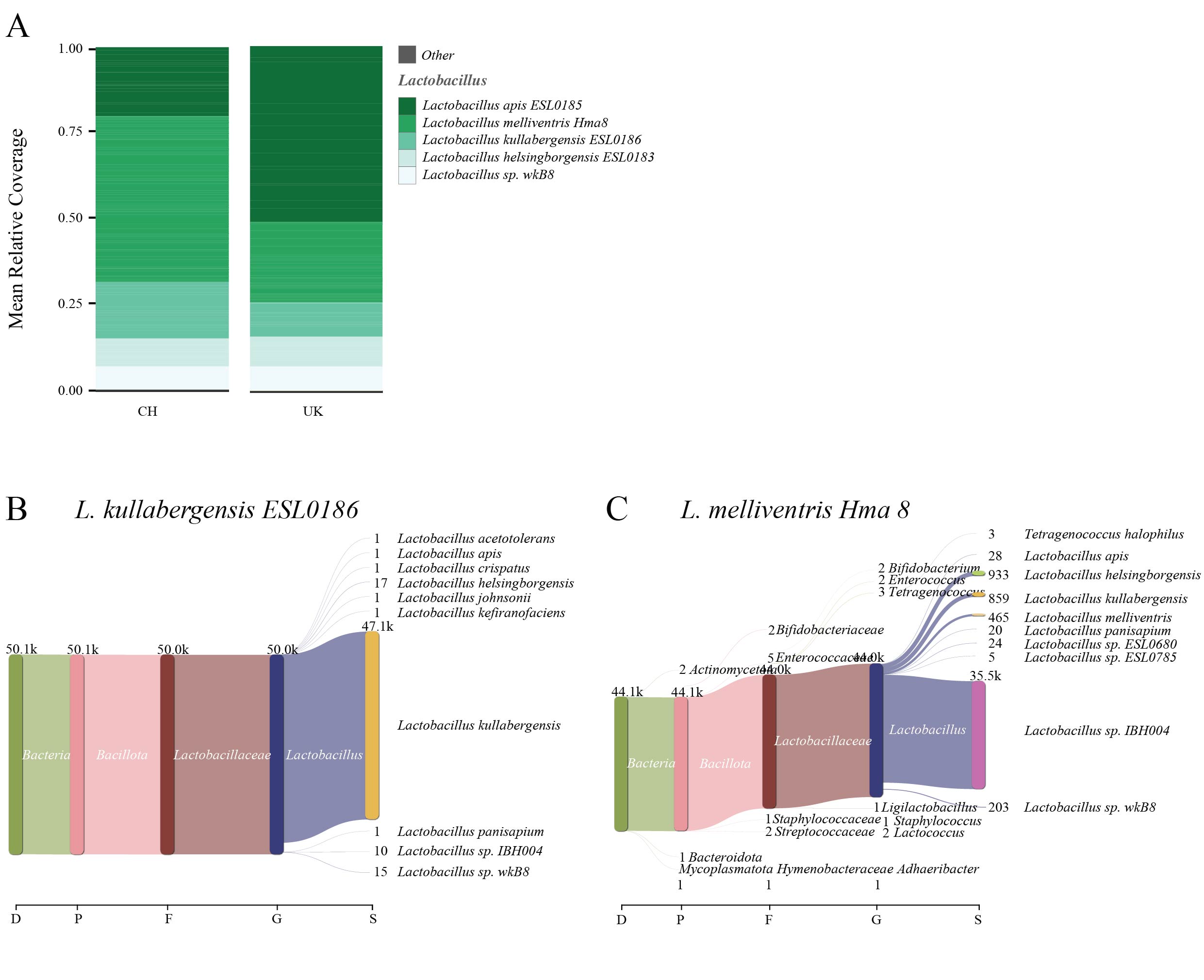

### Fig.S4.jpg

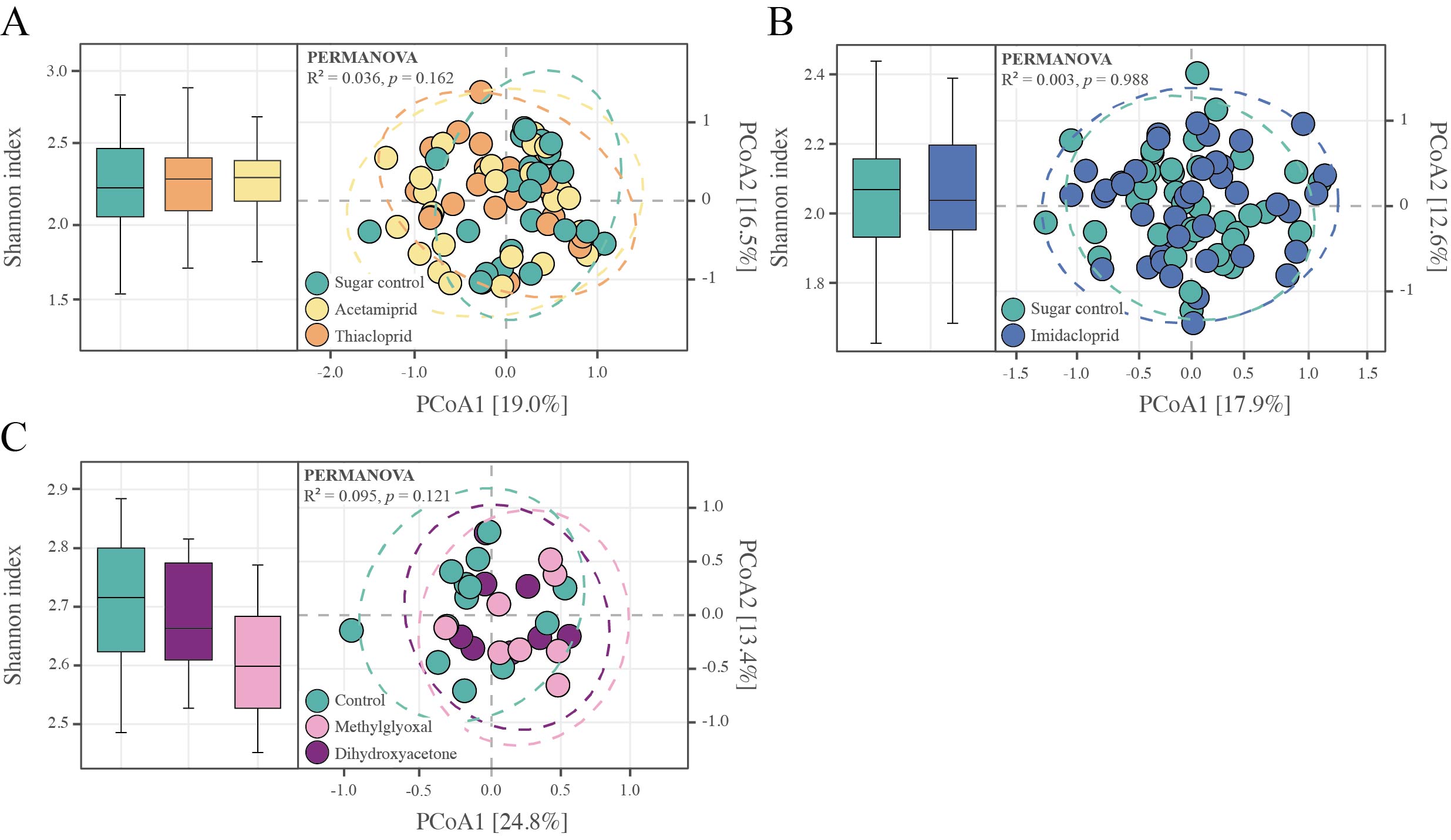

### Fig.S5.jpg

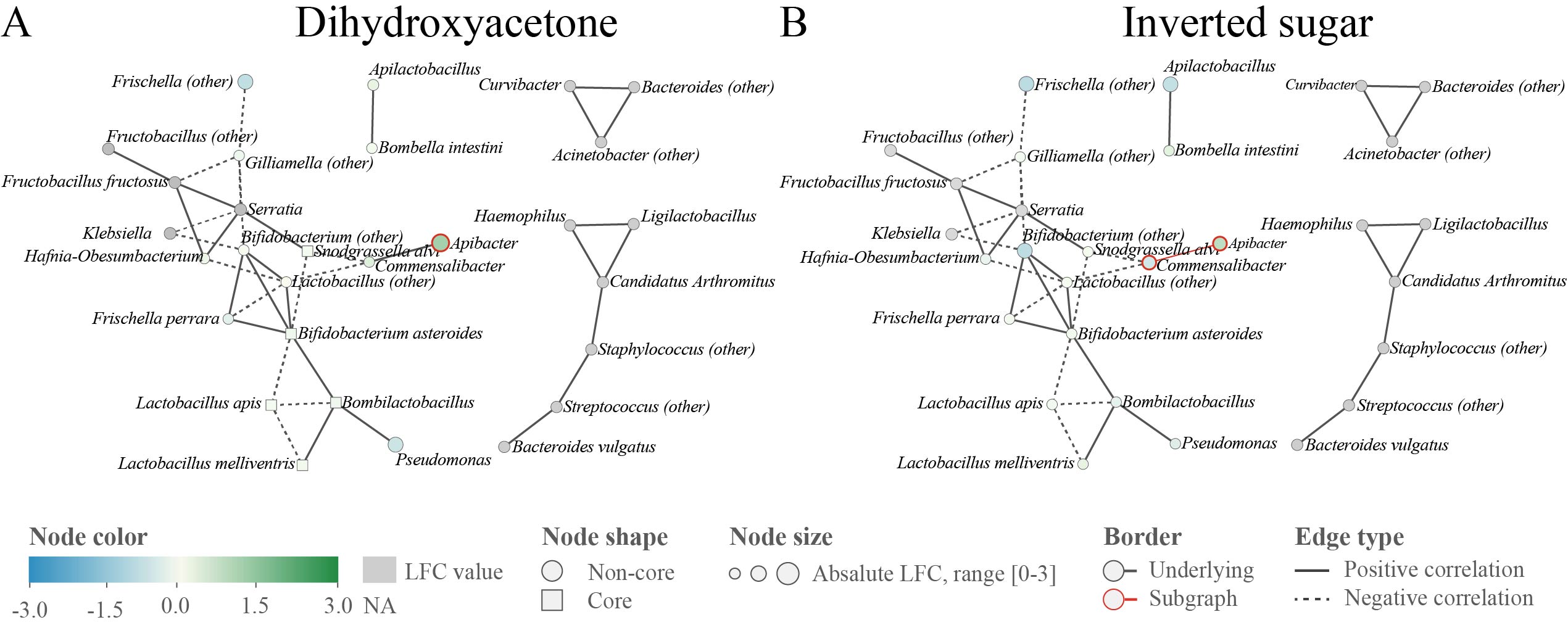
